# Uncovering and characterizing splice variants associated with survival in lung cancer patients

**DOI:** 10.1101/623876

**Authors:** Sean West, Sushil Kumar, Surinder K. Batra, Hesham Ali, Dario Ghersi

## Abstract

Splice variants have been shown to play an important role in tumor initiation and progression and can serve as novel cancer biomarkers. However, the clinical importance of individual splice variants and the mechanisms by which they can perturb cellular functions are still poorly understood. To address these issues, we developed an efficient and robust computational method to: (1) identify splice variants that are associated with patient survival in a statistically significant manner; and (2) predict rewired protein-protein interactions that may result from altered patterns of expression of such variants. We applied our method to the lung adenocarcinoma dataset from TCGA and identified splice variants that are significantly associated with patient survival and can alter protein-protein interactions. Among these variants, several are implicated in DNA repair through homologous recombination. To computationally validate our findings, we characterized the mutational signatures in patients, grouped by low and high expression of a splice variant associated with patient survival and involved in DNA repair. The results of the mutational signature analysis are in agreement with the molecular mechanism suggested by our method. To the best of our knowledge, this is the first attempt to build a computational approach to systematically identify splice variants associated with patient survival that can also generate experimentally testable, mechanistic hypotheses. Code for identifying survival-significant splice variants using the Null Empirically Estimated P-value method can be found at https://github.com/thecodingdoc/neep. Code for construction of Multi-Granularity Graphs to discover potential rewired protein interactions can be found at https://github.com/scwest/SINBAD. Presentation slides are found at https://github.com/scwest/RECOMB-CBB_2019_NEEP.

**Author summary:** In spite of many recent breakthroughs, there is still a pressing need for better ways to diagnose and treat cancer in ways that are specific to the unique biology of the disease. Novel computational methods applied to large-scale datasets can help us reach this goal more effectively. In this work we shed light on a still poorly understood biological process that is often aberrant in cancer and that can lead to tumor formation, progression, and invasion. This mechanism is alternative splicing and is the ability of one gene to code for many different variants with distinct functions. We developed a fast and statistically robust approach to identify splice variants that are significantly associated with patient survival. Then, we computationally characterized the protein products of these splice variants by identifying potential losses and gains of protein interactions that could explain their biological role in cancer. We applied our method to a lung adenocarcinoma dataset and identified several splice variants associated with patient survival that lose biologically important interactions. We conducted case studies and computationally validated some of our results by finding mutation signatures that support the molecular mechanism suggested by our method.

## Introduction

Large-scale cancer sequencing initiatives have opened up a window into the genome of individual cancers, offering unprecedented opportunities for studying the functional consequences of molecular alterations in human cancers [1]. However, despite new evidence highlighting the importance of cancer-specific splice variants in tumor initiation and progression, the role played by splice variants in the biology of human cancers and patient survival is still poorly understood.

To address this need, we developed a novel, statistically robust, multi-granular method to identify splice variants that are significantly associated with cancer patient survival. After clinically important splice variants are detected, our approach identifies variant-specific losses and gains of protein domains that can potentially rewire the cellular interactome, enabling a mechanistic interpretation of the role played by these splice variants. We applied this approach to Lung Adenocarcinoma (LUAD) using data from The Cancer Genome Atlas (TCGA), based on the clinical importance of the disease, the large number of observed point mutations that can lead to aberrant splicing, and the spread in patient survival time.

With lung cancer as the leading cause of cancer-related deaths [2], novel treatments are constantly developed to improve patient survival while reducing the toxicity and side-effects of conventional chemotherapy. In this regard, molecular targeted therapy is arguably one of the most investigated new paradigms in oncology. The general strategy in targeted therapy is to use small molecules to inhibit specific pathways important for cancer cell survival and invasiveness. [3]. However, for molecular targeted therapy to be successful, it is important to identify highly specific molecular mechanisms that lead to tumor survival and metastasis. For this reason, cancer-specific alternative splicing has received increasing attention for its role in tumor formation [4, 5], and methods to target splicing in cancer have recently been proposed [6] using RNAseq data.

While some argue in favor of the robustness of gene level analysis of RNAseq data [7], splice variant analysis offers unique information [8]. Sebestyén et al. developed a computational method to identify splicing signatures in cancers using TCGA data, showing that splicing signatures can be used to distinguish tumor from normal samples [9]. Yang and co-workers developed a web server called ISOexpresso to identify predominantly expressed isoforms in TCGA data [10]. Recent work by Climente-González has investigated the potential functional consequences of splice variants in TCGA datasets, highlighting Gene Ontology terms that are enriched in isoform switches and protein interactions that might be lost or gained [11]. We suggest that more detailed understanding of the influence of transcriptional expression on disease mechanisms in lung cancer may be gleaned from granularity-aware integration of RNASeq data with other sources of domain knowledge.

The strategy we propose here is to combine established RNAseq processing methods with patient survival analysis at the gene and splice variant level. Exon-skipping and splice variant level survival analysis have been shown to outperform gene level analysis at predicting patient survival [12]. Many methods for integration of RNAseq data exist and usually occur during downstream processing [13]. However, to the best of our knowledge this is the first attempt at building a high-throughput approach for survival analysis at the gene or splice variant level for early integration with domain sources.

Another important requirement for confident knowledge extraction is that of robustness, defined as stability of the results with respect to small changes in the parameters of the method. Non-parametric survival analysis requires a threshold that splits patients into subpopulations (e.g. an expression threshold). Previously, high-throughput survival analysis using arbitrary thresholds were shown to be sensitive to changes in patient groups [14]. One known approach for increasing this robustness is to find the minimum *p*-value by sampling multiple thresholds. However, this produces *p*-values that are non-uniform and skewed. Lausen and Schumacher examined this problem for the logrank test, developing an equation for estimating the corrected *p*-values [15]. Yet, this type of correction method was not developed for high-throughput analysis, where multiple hypothesis testing is required and is dependent on very small changes of individual *p*-values. To overcome these issues, we developed a reliable and high-throughput survival analysis method called NEEP (null empirically estimated *p*-values). This method requires no distribution assumptions, guarantees statistically valid *p*-values, and may be implemented in a high-throughput way.

We applied this method to the LUAD RNAseq dataset from TCGA to conduct survival analysis at the splice variant level, identifying variants that are significantly associated with patient survival. Then, we integrated these results with protein-domain annotation and protein-protein interaction data to provide a mechanistic interpretation for the potential role played by these clinically relevant splice variants in tumor formation and progression. More specifically, we identified variants that lose or gain protein domains involved in mediating protein-protein interactions. We refer to this integration approach as a multi-granularity graph (MGG). Fig. 1 shows the overall workflow of the method.

**Fig 1.**
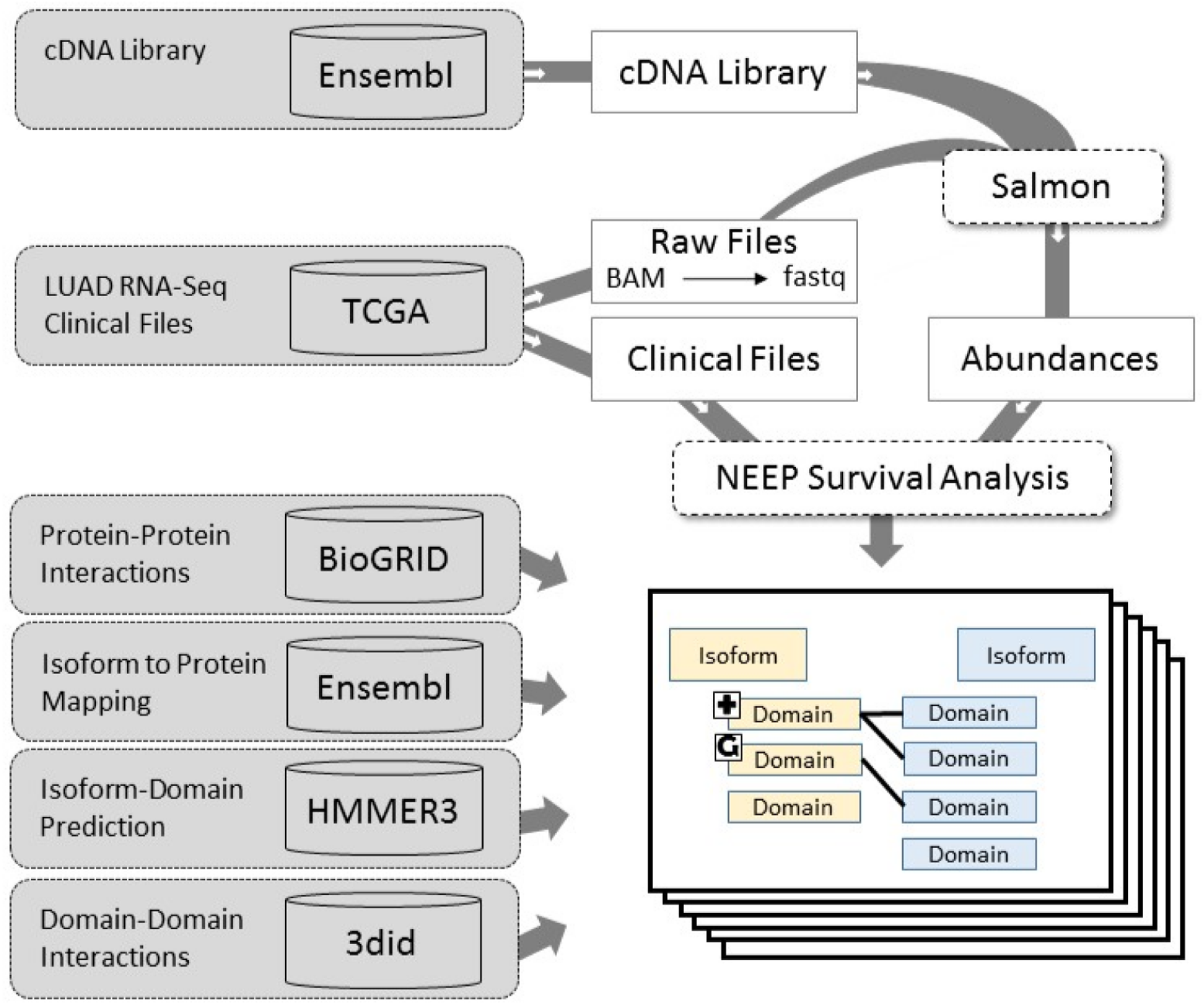
Workflow for generating multi-granularity graphs (MGGs). Using the NEEP method significant splice variants were calculated, utilizing clinical data, the complementary-DNA reference library, and RNA-Seq data from Ensembl and TCGA. Multi-granular graphs were constructed for known protein-protein interactions from BioGRID, mapped from Ensembl to at least one significant splice variant. Domains were predicted for each splice variant using HMMER3 and Pfam. Gained and ghost domains were identified in splice variants associated with survival. Final MGGs represent potential lost or gained protein interactions associated with patient survival.

In the absence of a gold standard, we conducted case study analysis with literature review to assess the biological plausibility of the inferred graphs. In summary, our main contributions are the following: (1) we use the splice variant granularity level as a way to identify changes in protein mechanisms that can affect patient survival; (2) we present a novel, computationally efficient, and robust method to perform high-throughput survival analysis; and (3) we integrate multiple data types for inferring biologically plausible and potentially actionable molecular targets in lung cancer.

## Results

### NEEP identifies splice variants significantly associated with patient survival

One of the contributions of this work is the development of a statistically robust and computationally efficient method to identify optimal expression thresholds that yield minimum *p*-values when performing survival analysis over a large number of transcripts. The efficiency of the NEEP method is due to the fact that we can estimate empirical *p*-values for all transcripts by computing only one null distribution, as opposed to having to perform sample permutations for every transcript (as discussed in detail in Materials and Methods). To visually show this, we plot the *p*-value distributions of 100 randomized gene expression patterns of size 1000, against 20 simulated null distributions of size 1000 obtained by shuffling samples (Fig 2A). The overlap between these distributions visually confirms that the same null distribution can be utilized across all expression patterns without the need for expensive sample permutations for every transcript.

**Fig 2.**
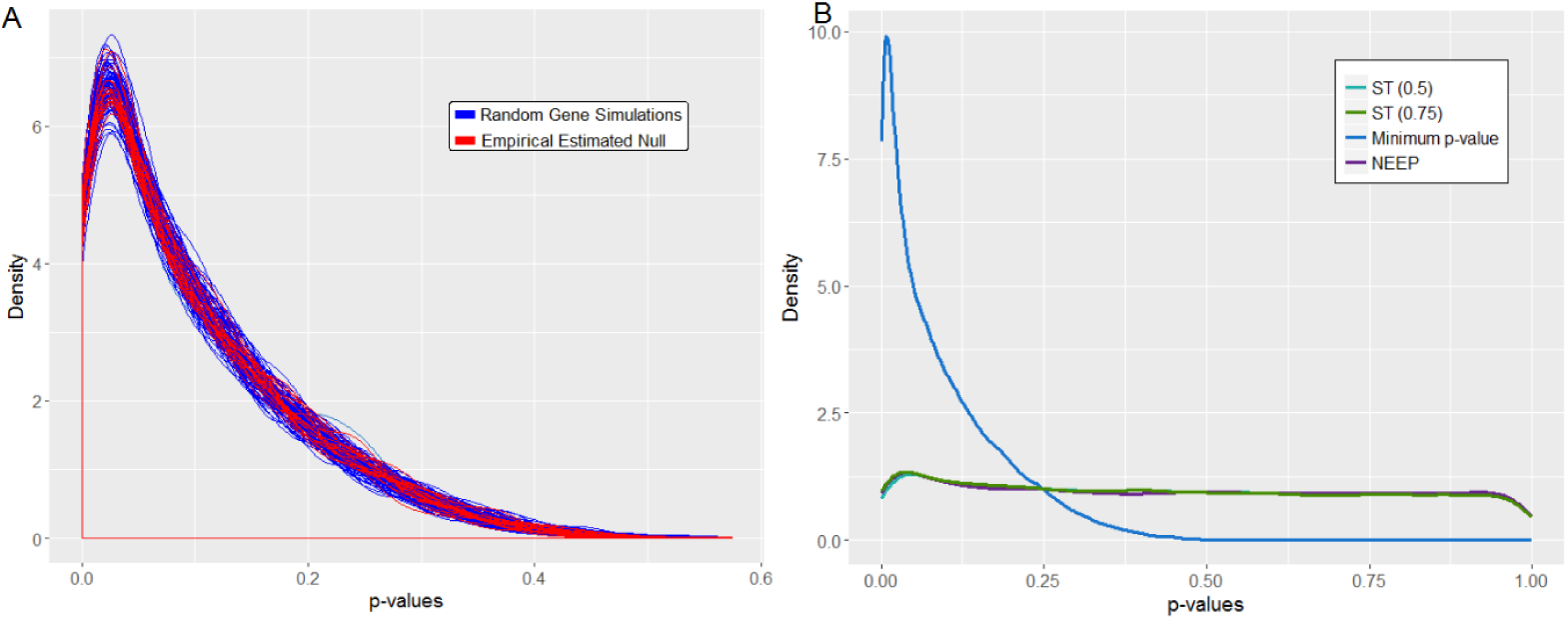
NEEP yields uniformly distributed *p*-values. (A) The *p*-value distribution is shown for the minimum *p*-value method and NEEP. An ideal statistical test produces *p*-values that are uniform under the null. (B) Density line plots were constructed for each of 100 simulations of 1000 random abundance patterns (blue) and 20 sets of 1000 values chosen at random from the initial 1,000,000 null distribution (red). The overlap between the two groups of density plots empirically confirms that the null distribution is the same for all expression patterns, given identical clinical data.

We ran NEEP using 10 randomly generated null distributions of 1,000,000 isoform simulations. In all 10 repetitions, there were 181 NEEP significant (FDR < 0.1) splice variants from the LUAD dataset. Since the order of splice variants in NEEP does not depend on the null but only their statistical significance does, these 181 splice variants were identical each time. The mean hazard ratio of the significant splice variants was 3.122 and the minimum was 2.18.

The distribution of all the *p*-values is nearly uniform, as can be seen in Fig 2B. The small deviation of low *p*-values from the uniform distribution is due (as expected) to the significant splice variants.

### Case study of multi-granular graphs linked to DNA repair

After identifying splice variants significantly associated with survival, we followed the procedure described in Materials and Methods and summarized in Fig. 1 to generate multi-granular graphs (MGGs). Briefly, MGGs show splice variant specific protein-protein interactions that might be gained or lost as a result of gains or losses of protein domains in splice variants significantly associated with patient survival.

After constructing the MGGs using the NEEP significant splice variants, we obtained 50 MGGs (the complete list is available in S3 File). We first checked whether the MGGs we obtained were representative of the full set of significant splice variants identified by NEEP. To do that, we performed Gene Ontology enrichment analysis on NEEP significant genes, NEEP significant isoforms, and genes represented in MGGs, as discussed in S1 File. All GO enrichment term groups obtained at the gene or isoform levels were also present in the MGGs, as shown in S1 Fig.

Using these MGGs and information available from the literature, we investigated possible mechanisms that can explain why these splice variants are correlated with survival of lung cancer patients. Here we discuss three case studies linked to DNA repair. Additional MGGs are discussed in the supplemental materials S2 File. All discussed MGGs are visualized in Fig 3.

**Fig 3.**
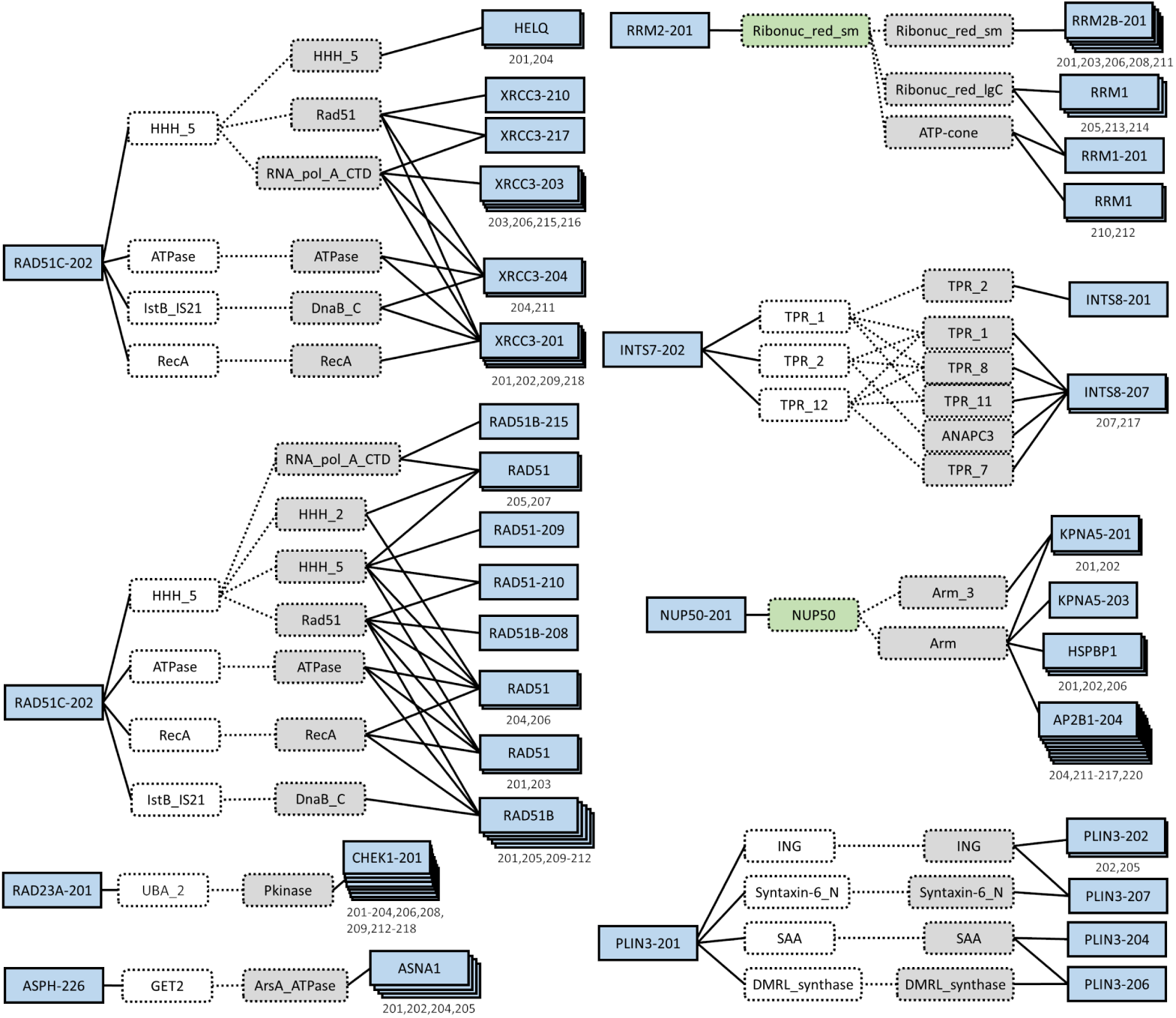
Multi-granularity graphs. Splice variants are in blue solid boxes and domains are in dotted boxes. The significant splice variant (left box) is linked to its gained (green) and ghost (white) domains. Domains in gray belong to the non-significant splice variant (right box). Splice variants of the same gene that have identical domain connections in the MGG are stacked and their identifiers are listed below the boxes. RAD51C-202 has two MGGs which are separated to simplify visualization.

### RAD51C-202 loses ability to bind to the Holliday junction, XRCC3, and HELQ

NEEP suggests that the increased expression of RAD51C-202 is associated with lower patient survival. Further, the interaction between RAD51C-202 and XRCC3 is prevented because of several missing domains. Upon deeper inspection using RCBP PDB and Ensembl, we see that RAD51C-202 is missing two exons that are present in the remaining RAD51C isoforms. The first exon allows RAD51C to bind to the Holliday junction, a relationship missed by the MGG. The second exon contains the domains which allows RAD51C to bind to XRCC3. RAD51C and XRCC3 have long been known to form a complex (CX3 complex) which regulates genetic recombination and DNA repair [31]. Dysruption of this complex (perhaps via increased expression of RAD51C-202) may promote cancer in two ways. First, loss of RAD51C has been implicated in tumorigenesis [32]. Second, it was suggested that the CX3 complex increases mitochondrial stability, reducing the risk of cancer [33]. In addition, HELQ interacts with RAD51C within the BCDX2 complex [34]. Disruption of the BCDX2 complex increases cellular susceptibility to agents that induce DNA-interstrand crosslinks [35]. Thus, the additional transcription of the RAD51C-202 isoform which is missing the domain necessary to bind to HELQ may result in DNA-interstrand crosslinks.

### RAD23A-201 loses interaction with CHEK1 proteins

The RAD23A-201 isoform does not include the UBA 2 domain, which is found in all of the other RAD23A isoforms. The survival analysis indicates that a high expression of RAD23A-201 is associated with lower patient survival. RAD23A operates by bringing proteins marked for ubiquitination to the proteasome, avoiding its own degradation. Without UBE 2 RAD23A is ubiquitinized, preventing its role in nucleotide excision DNA repair and protein degredation [36]. Further, without the UBA 2 domain, the PPI between RAD23A and CHEK1 via the DDI between UBE 2 and Pkinase will not occur. It may be possible that RAD23A phosphorylates CHEK1 since its cousin, RAD17, is a known phosphorylator of CHEK1 [37]. However, this is unlikely since RAD23A is known to function primarily through its association with the proteasome [38]. The inhibition of the PPI with CHEK1 may prevent degredation of CHEK1. Under several circumstances CHEK1 promotes tumor growth and increases therapy resistance [37]. Thus the reduction of RAD23A with longevity could inhibit CHEK1 degredation and allow promotion of tumor growth and therapy resistance.

### INTS7-202 has altered structure and number of TPR domains

INTS proteins serve as members of the Integrator complex (INT). The roles of this complex are diverse but suggestive of potential roles in cancer, such as mediation of RNA polymerase II or DNA damage response [39]. Yet the INT is ‘modular’, containing different subunits of INTS genes and causing different effects on the role of INT [40]. The MGG model suggests that the interaction between INTS7 and INTS8 may be effected by the loss of a few TPR domains in INTS7-202. The TPR domains are common PPI domains which allow the construction of many protein complexes, including INT. Losses or disruptions of these domains in INTS7-202 may cause instability and prevent the formation of INT. Alternatively, given that many other TPR domains are uneffected by the alternative splicing of INTS7-202, INT may still form but function differently. In addition, INTS proteins also function outside of the INT complex [39].

### RAD51C-202 expression is linked to a characteristic mutational signature and lower patient survival

The biological models discussed above suggest a role played by these splice variants in DNA repair. Alterations in DNA repair mechanisms result in characteristic mutations signatures. We hypothesized that patients in the RAD51C-202 low expression and high expression groups might have different mutation signatures reflecting different abilities to perform DNA repair via homologous recombination.

As shown in Table 1, RAD51C-202 is associated with COSMIC mutation signatures 2, 3, 11, 12, 16, 17, and 23. Of particular interest, signature 3 (Fig 4) is reported by COSMIC as associated with failure in DNA double-strand breaks in homologous recombination. Indeed, the results of the Mood’s test indicate that signature 3 is more prevalent in patients with high expression of RAD51C-202 than in patients with low expression (*p* < 0.0232, Fig 4). RAD51C-202 had the highest association with COSMIC signature 2, which is known to be caused by AID/APOBEC activity. However, we could not find more information from the literature regarding this particular signature.

**Table 1.** COSMIC mutation signatures associated with RAD51C-202 expression.

|  | High - Low (Mean Contribution) | Description from COSMIC |
| --- | --- | --- |
| Signature2 | 11.032 | Activity of AID/APOBEC deaminases |
| Signature3 | 3.864 | DNA double-strand break-repair by homologous recombination |
| Signature11 | 0.809 | Similar mutation pattern to alkylating agents |
| Signature12 | 2.323 | Unknown |
| Signature16 | 3.194 | Unknown |
| Signature17 | -0.016 | Unknown |
| Signature23 | 0.533 | Unknown |

The difference between the mean contributions of *low* from *high* RAD51C-202 expression groups to each significant mutation 181 signature is reported. Signature 4, associated with smoking, is not significant. 182

**Fig 4.**
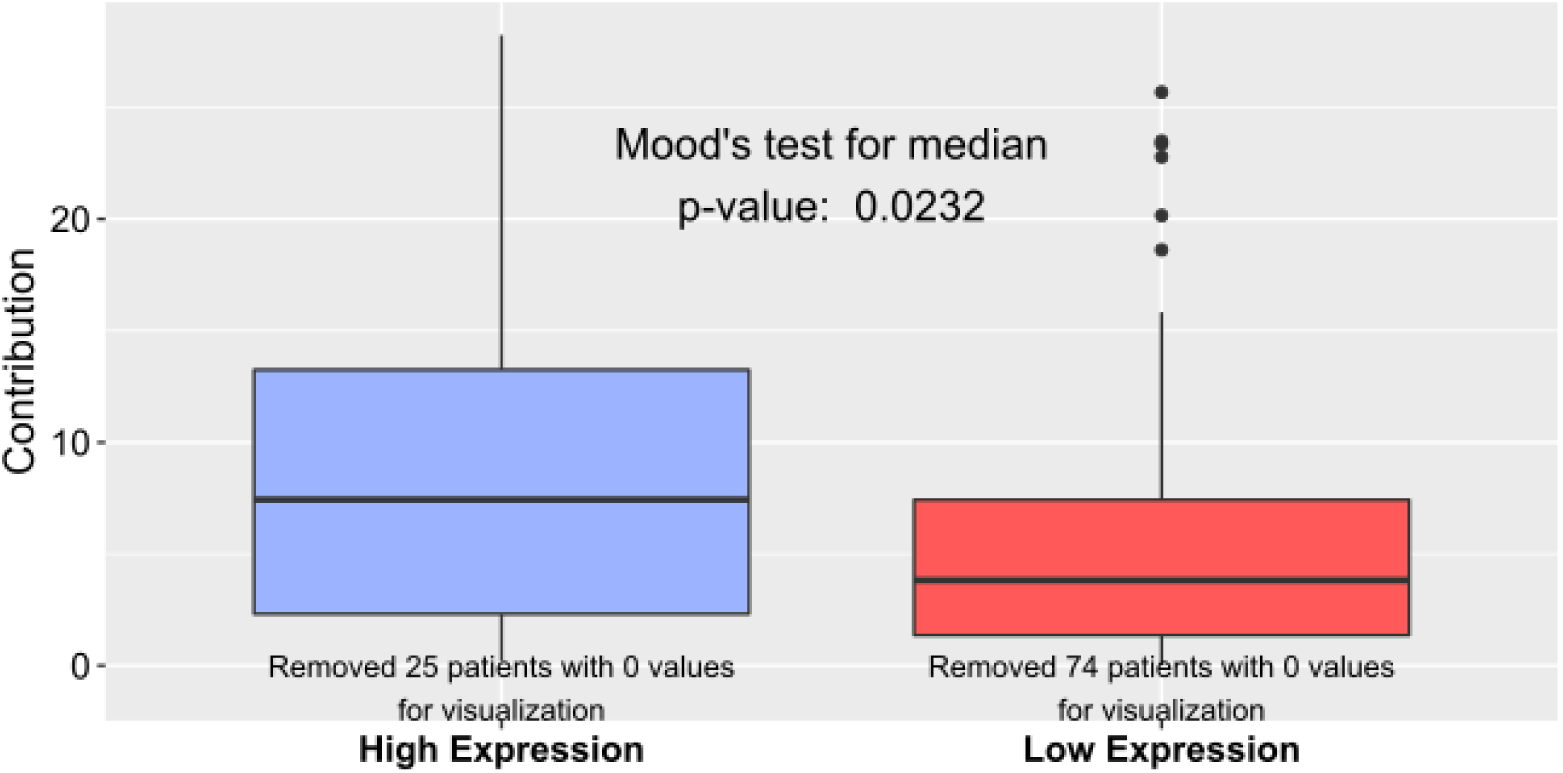
Mutation signatures associated with RAD51C and smoking effects. (A) The mean contribution of each patient in the high RAD51C-202 expression group was subtracted from the low group. A positive number indicates more contribution of the COSMIC mutation signature to the patients in the high expression group. Listed signatures have a significant Mood’s test p-value for RAD51C-202 expression. (B) Potential smoking confounding variables were separated by RAD51C-202 expression and tested according to their data type. (C) Box-plots were generated for the signature-3 contribution to patients in low (red) and high (blue) expression of RAD51C-202.

Since smoking is commonly associated with lung cancer and may cause particular mutation patterns, the relationship between RAD51C-202 with signature 3 could be confounded by smoking. However, smoking does not appear to be a confounding factor in RAD51C-202 association with COSMIC gene signatures. As can be seen in Table 2, no smoking variable was significantly different between patients with low and high expression of RAD51C-202. Further, signature 4 contribution (found in smokers) was not significantly associated with RAD51C-202 abundance. In addition, survival does not appear to be a confounding factor. The survival Cox-PH *p*-value for RAD51C-202 was significant with and without using signature 3 contribution as a confounding factor (*p*-values: 0.010 and 0.036). The proportion of NEEP significant isoforms that were associated with signature 3 was insignificant (*p*-value = 0.6403).

**Table 2.** Statistical test results comparing smoking variables to RAD51C-202 Expression

| Smoking Variable | Data Type | Statistical Test | $p$-value |
| --- | --- | --- | --- |
| Cigarettes Per Day | Metric | Welch Two Sample T-test | 0.086 |
| Years Smoked | Integer | Wilcoxon Rank Sum Test | 0.098 |
| Is Smoker | Binary | 2-Sample Test for Equal Proportions | 0.383 |

### Robustness of NEEP to changes in patient population

We conducted 100 simulations by removing 5% and 10% of the patients at random prior to running NEEP. We compared the rank of the significant isoforms observed in the full dataset to their rank in the simulations. Fig 5A shows the range of the middle 50% ranks from the simulations, and it also plots the original, observed values. Of the 181 observed significant splice variants, few had ranks greater than 400 out of 69,561. Further, splice variants with lower observed rank had lower variance in the simulations. In addition and as expected, ranks in the 5% simulations were much lower than the ranks in the 10% simulations. Fig 5B is the density plot of the observed ranks paired with the 10% simulation ranks. The density nearly follows the *y* = *x* line, where the observed rank is the same as the simulated rank. Once again we see that splice variants with low observed rank have lower simulation variance than variants of higher rank. Mean Spearman rank correlations between the entire set of observed splice variants and the 5% and 10% simulation rounds were 0.861 and 0.931, respectively. Overall, these results suggest that the NEEP method is fairly robust to removal of patient samples.

**Fig 5.**
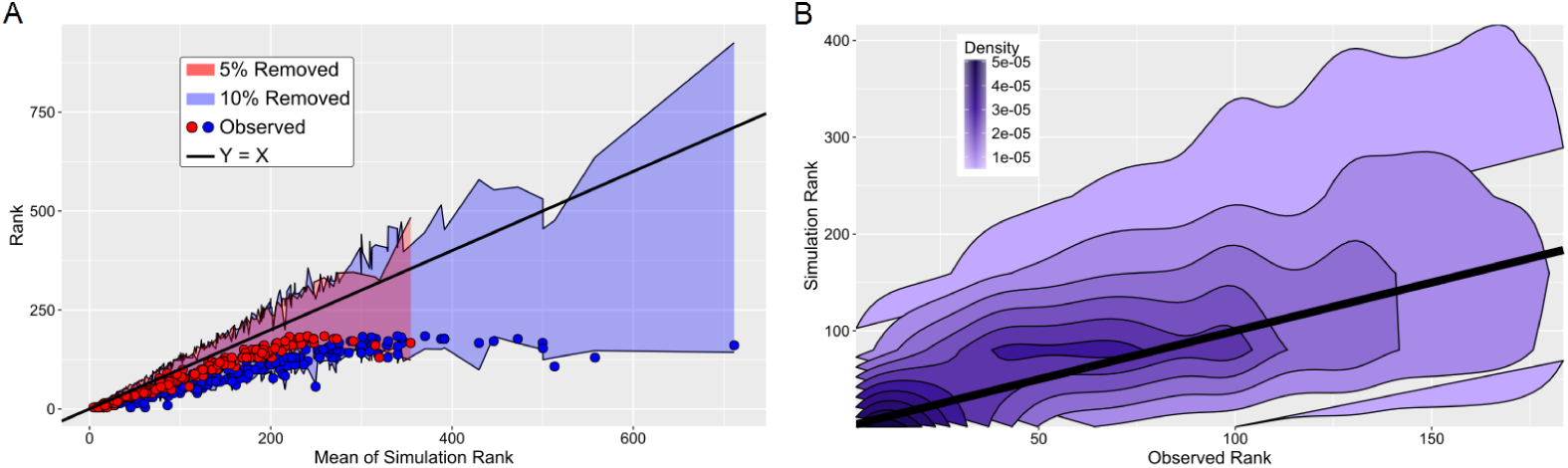
Robustness of single threshold methods and NEEP. (A) The central 50 ranks for the 100 simulations of each of the 181 significant splice variants are plotted as a shaded range according to their mean simulated rank, separately for the 5% simulations (red) and the 10% simulations. Each of these ranges correspond to an observed (non-simulated) rank, which is plotted as a dot along the same x-axis. The 5% simulation ranges have lower means of simulation rank than the 10% simulation ranges; thus the red dots are closer to the identity line. The maximum mean of the simulation rank is lower for the 5% simulations; thus the red shade ends much sooner than the blue shade. (B) The density plot of the observed vs. simulation pairs for the 181 significant splice variants.

## Discussion

An exploratory survival analysis must be high-throughput to span all genes/splice variants (in this study, 21,100 genes and 69,561 splice variants). Cox regression and Kaplan-Meier (KM) curves are two options. We chose to avoid Cox regression for two reasons. First, we violate the Cox regression assumption that the mechanism of censoring is not related to patient survival, as patients who survive to the end of the study will have a higher survival rate than the overall population at the outset. As discussed in the Methods, we can choose a threshold range which prevents loss of statistical power due to this phenomena on the KM test. Second, Cox regression firmly assumes the proportional hazards assumption, where the assumption is looser in KM analysis. The effect of many splice variants on survival is not constant over time, resulting in varying hazard rates. Driver genes are just one example that violate this assumption.

The hypothesis of the KM logrank test is also better suited to this problem than Cox regression: Cox regression tests against the null that the variability of the survival cannot be explained by the variability of the splice variant, whereas the KM logrank test compares *low* against *high* expression. Therefore, each KM hypothesis is associated with the maximum difference between two survival curves, and exact delineations of *low* and *high* expression are extracted.

A current limitation of our approach is that the multi-granularity graphs only highlight splice variants that can gain or lose protein-protein interactions as a result of gains/losses of protein domains. Protein-DNA, protein-RNA, and protein-ligand interactions are not considered. We plan to extend our work in this direction in the near future.

In conclusion, we presented NEEP, a novel computational method to systematically identify splice variants significantly associated with patient survival. An important advantage of NEEP over the existing methods is that the optimal expression threshold to separate patients into better vs. worse survival groups is automatically detected by the method, and the resulting *p*-values are uniformly distributed under the null hypothesis, allowing for proper multiple hypothesis correction.

By applying our method to the TCGA lung adenocarcinoma dataset, we identified several splice variants that significantly correlate with patient survival. We then built multiparted, multi-granular graphs (splice variant-domain-domain-splice variant) that allowed us to identify putative losses and gains of protein-protein interactions associated with these splice variants. Several of these variants turned out to be implicated in DNA repair through homologous recombination. To computationally verify this, we performed a mutational signature analysis that supported the molecular mechanism suggested by the multi-granular graphs.

## Materials and methods

### Data retrieval and transcriptome profiling

We downloaded the TCGA Lung Adenocarcinoma (LUAD) dataset from the Cancer Genomics Hub (CGHub). Of the 515 patients with RNA-Seq data, 75 were removed due to clinical annotations suggesting possible confounding factors like “Prior malignancy” or “Synchronous malignancy”. Eight patients were removed due to missing survival data, resulting in 432 total patients. To reconstruct splice variant abundance estimates, the TCGA BAM files were reverted back to fastq format using SAMtools [16].

We used Salmon [17] with default parameters to estimate transcript-level abundances based on the Ensembl 91 cDNA library. Non-alignment methods such as Salmon offer accurate abundance estimates at the splice variant level [18]. In addition, Salmon performed best during a comparison of alignment-free methods [19]. Using Ensembl gene/splice variant mapping, we summed splice variant abundances to estimate gene abundance. TPM values were used to represent the gene/splice variant abundances. We note that since non-parametric survival analysis utilizes two Kaplan-Meier curves for each gene/splice variant (one for *high* abundance and one for *low* abundance), the choice of using counts, FPKM or TPM will result in identical logrank *p*-values.

### High-throughput survival analysis

We performed non-parametric survival analysis at the gene and splice variant level using a previously published approach (“minimum *p*-value”) and our new NEEP (“null empirically estimated *p*-values”) method, as described below. The clinical files containing survival data were obtained from TCGA. The *days*_*to*_*death* and *days*_*to*_*last*_*followup* were used as uncensored and censored survival values, respectively.

#### Minimum *p*-value approach

Choosing the threshold which maximizes a test statistic avoids the choice of an arbitrary threshold, while increasing reliability. Minimum *p*-values were calculated for each gene and splice variant. The logrank test was performed on each possible split between high and low abundance within a threshold range of 0.15 to 0.85. This method results in a distribution of *p*-values that is non-uniform and skewed to the left.

#### Empirical estimation of *p*-values

To overcome the limitations of choosing a single threshold (arbitrary threshold) and minimum *p*-value methods (non-uniform *p*-values under the null hypothesis), we developed a new approach resulting in null empirically estimated *p*-values (NEEP). The method works by estimating the null distribution of the minimum *p*-value approach, so that *p*-values under the null hypothesis are uniformly distributed, allowing for *p*-value adjustment.

When using the logrank test with a single expression threshold *t*, the set of all possible *p*-values is finite, and it corresponds to all possible patient partitions into the *low* and *high* expression groups. By extension, the null distribution of the minimum *p*-value method is defined as the entire discrete distribution of possible *p*-values across all values of *t*. The exact number of possible *p*-values can be calculated as a function of *s* (the number of samples), *l* (the minimum percentile threshold), and *h* (the maximum percentile threshold), according to the following equation:

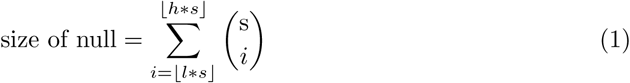

where ⌊*x*⌋ is the greatest integer less than or equal to *x*. To determine a viable range of thresholds, we first chose minimum power and effect size. Given a sample size of 432 patients with 151 recorded death events and an alpha of 0.1, to obtain a hazard ratio greater than 2 (or less than 0.5) and a minimum power of 0.75, the survival curve with fewest patients must have at least 62 members. This corresponds to a minimum percentile threshold of 0.143, which we round in this study to 0.15 for simplicity. So, in this study we set *l* to 0.15 and *h* to 0.85.

Considering the size of *s*, enumerating all possible *p*-values is computationally unfeasible. Therefore, we built a null distribution with 1,000,000 Monte Carlo simulations performed by randomly partitioning samples into the two groups using all possible thresholds and extracting the smallest logrank test *p*-value. We estimated *p*-values as the fraction of simulated values less than or equal to the minimum *p*-values obtained from the actual data. As expected, the NEEP procedure yielded uniformly distributed *p*-values. Benjamini-Hochberg adjustment was conducted on the NEEP *p*-values, and adjusted values less than 0.1 were considered significant.

It is noteworthy to mention that individual expression values do not impact the KM survival curve as long as they result in the same split between high and low expression. Therefore, the NEEP procedure needs to be carried out only once for the whole set of genes/splice variants, provided the fraction of patients expressing the gene/splice variant is as large as or greater than 1 - *l*, the minimum percentile threshold. In our dataset, this criterion was met by 21,100 genes and 69,561 splice variants. Thus, we empirically estimated the *p*-values of these genes/splice variants using a single sampled null distribution. While effect sizes of the logrank test are not as naturally interpretable as many other statistical tests, we calculated the hazard rate ratio of *high* over *low* expression curves as well as the mortality rates at 1, 2, and 5 years. Code for constructing NEEP values from clinical and expression data and calculating effect size statistics is available at https://github.com/thecodingdoc/neep.

### Robustness

We measured robustness of the NEEP method by examining sensitivity of significant splice variants to changes in the set of patients. We conducted two rounds of 100 simulations, where 5% and 10% of patients were randomly removed, respectively. Since the number of patients in each simulation is then compared to the actual values, we expect the adjusted NEEP *p*-value change is due to a smaller sample size. So we compared ranks instead of *p*-values to isolate the sensitivity of NEEP *p*-values to resampling. We compared ranks of the significant isoforms in the full dataset against their rank in the simulations.

### Multi-granular graphs

We utilized data across multiple granularities to construct plausible graphs for the association between splice variant expression and lung cancer survival. Specifically, we used protein-protein interactions (PPI) from BioGRID [20], domain-domain interactions (DDI) from 3did [21], splice variant domain prediction using HMMER3.1 [22], and the survival analysis significance values using NEEP, with the goal of identifying cancer-specific changes in PPI interactions.

The general structure of these multi-granular graphs (MGG) can be seen in Fig 1. The outcome of the workflow consists of quadripartite networks, connecting splice variants with their potential interaction partners via their domain-domain interaction, as *variant-domain-domain-variant* paths.

To build these graphs, we extracted protein-protein interactions from BioGRID v3.4.156 [20], retaining only physical interactions in human, resulting in 377,228 relationships. Of these interactions, 348,609 involved proteins with an active Ensembl ID. Because PPIs are reported at the gene product level, we expanded the interactions to include all possible variant-variant interactions (73,588,825). To identify graphs associated with survival, the left-most splice variant is required to be both survival-significant (NEEP FDR < 0.1) and protein-coding, leaving 655,407 unique possible interactions.

We identify two domain patterns of interest, “ghost” and “gained”. Ghost domains are those that are present in survival-insignificant splice variants within the same gene, but are not found in the survival-significant splice variant. Gained domains are found in the survival-insignificant splice variants but present in the survival-significant splice variants. By incorporating these gained and ghost domains direct into their PPI context, we can consider MGGs as potential interaction changes caused by domain exclusion or inclusion. Genes without multiple splice variants were removed from further analysis. We downloaded the 11,200 domain-domain interactions (DDI) from 3did [21], where inclusion requires a known protein-protein structure to support the interaction. Variant-domain membership was determined using HMMER3.1 [23] with default parameters and the Pfam database [24]. After removal of interactions without DDI support, 5912 interactions remained. Of these, only 1208 had gained or ghost domains. There were 50 genes with an isoform significant for survival for these 1208 interactions. Code for constructing MGG interaction paths from variable data sources is available at https://github.com/scwest/SINBAD.

### Identification of significant COSMIC mutation signatures

To examine if the splice variant RAD51C-202 was associated with homologous recombination mutations, we utilized COSMIC gene signatures [25]. We used the optimal NEEP threshold of RAD51C-202 to separate the patients into a low expression (344 patients) and high expression (88 patients) group. Mutation data for each patient was downloaded from TCGA for all 4 variant callers: MuSE [26], MuTect [27], SomaticSniper [28], and VarScan [29]. Mutation profiles for each patient were constructed as described as the 6 substitution types along with the neighboring bases for a total of 96 mutation types using only mutations found by all four variant callers. The Catalogue of Somatic Mutations In Cancer (COSMIC) has identified a set of 30 commonly identified mutation signatures deconvolved from cancer patient mutation profiles [25]. We used MutationalPatterns [30] in R to find the linear contributions of each of the 30 signatures to a patient profile. We used Mood’s test for median to determine if the contributions were significantly different between low and high RAD51C-202 expression groups for each signature.

To check for possible confounding factors, we considered three smoking variables reported by TCGA and checked whether they were different between the low and high RAD51C-202 expression groups: *cigarettes per day* was tested using the Welch two sample t-test; *years smoked* was tested using the Wilcoxon rank sum test; and the binary *smoker* variable was tested using the two-sample test for equal proportions. In addition, we checked if survival confounded the relationship between RAD51C-202 and signature 3. Because the causality between mutations and survival must be directional, we conducted Cox-PH survival analysis of RAD51C-202 with and without the contribution of Signature 3 as a confounding variable. Further, Mood’s test was also conducted for the relationships between Signature 3 and the remaining splice variants with MGGs. The exact binomial test was used to determine if the proportion of splice variants significant for Signature 3 was greater than expected by chance.

## Supporting information

List of multi-granularity graphs.

Additional case studies of MGGs.

Enrichment analysis methods.

Enrichment term cluster membership across granularities.

## Supporting information

**S1 File. Enrichment analysis methods.**

**S1 Fig. Enrichment term cluster membership across granularities.** Counts of genes belonging to each enrichment cluster for the gene, isoform (splice variant), and MGG granularities are displayed as bar lengths. Missing bars does not indicate no membership, just insignificant enrichment of any term in the cluster.

**S2 File. Additional case studies of MGGs.**

**S3 File. List of multi-granularity graphs.** This table reports the entire list of MGGs for the TCGA LUAD dataset. The file does not contain the exact paths, only the members of each component of the MGG.

## Acknowledgments

This research was partly funded by a University of Nebraska Collaboration Initiative/System Science Seed Grant to SK, HA, and DG and by the NIH AA026428 R21 grant to SK.

