## Additional case studies of MGGs. for "Uncovering and characterizing splice variants associated with survival in lung cancer patients"

### Supplemental material: Additional case studies of MGGs Paper: Uncovering and characterizing splice variants associated with survival in lung cancer patients

Sean West[1], Sushil Kumar[2], Surinder K. Batra[2], Hesham Ali[1*], Dario Ghersi[1*]

**1** College of Information and Science Technology, University of Nebraska at Omaha, Omaha, NE, USA **2** Department of Biochemistry and Molecular Biology, University of Nebraska Medical Center, Omaha, NE, USA

*

#### Additional case studies of MGGs

##### PLIN3-201 cannot form homo-dimers

PLIN3-201 (TIP47) lacks several domains which are required for self-interaction. Also a higher expression of PLIN3-201 is associated with lower patient survival. At first glance, little research exists to connect the functionality of PLIN3 to cancer. However, down-regulation of PLIN3 has been shown to reduce the presence of lipid droplets [1]. Lipid droplet formation has been implicated in cancer cell proliferation [2]. Additionally, lipid droplet formation oscillations have been observed in lung cancer cells [3]. PLIN3-201 has a high ratio of expression over the other isoforms (0.660). Thus, differences in expression of this isoform may play a direct role in the biological process of lipid droplet formation. Active versions of PLIN3 have been observed forming trimers *in vitro* during migration of PLIN3 to the lipid droplet [4]. While little research has uncovered peptide configurations at the site of the PLIN3 interaction in lipid droplets, the MGG model suggests that formation of dimers or trimers may play an impactful role.

##### NUP50-201 gains interactions with KPNA5, HSPBP1, and AP2B1 proteins

The NUP50-201 isoform is the only isoform of NUP50 that contains a non-dysrupted NUP50 domain, after the removal of isoforms without expression in more than 66 patients. One role of NUP50 is in a nuclear import complex. Specifically, NUP50 forms an interaction with NUP153 known as the nuclear import complex [5]. Their association results from two interactions, one directly between NUP153 and one mediated by importin alpha [5]. The MGG suggests that the increase in the expression of NUP50-201, potentially the only isoform capable of interacting with KPNA5 (importin subunit alpha-6), is associated with a lower survival rate in lung cancer patients. Indeed, increased nuclear transport has been associated with multiple cancers [6] [7] [8]. Two alternative mechanisms are suggested in the MGG: with HSPBP1 and AP2B1. HSPBP1 contains the Arm domain which has specificity for the NUP50 domain in NUP50-201. HSPBP1 prevents ubiquitination of HSP70 [9]. It may be possible that HSPBP1 enhances NUP50 activity by increasing its lifespan. However, upon further inspection of BioGRID, this PPI received a very low interaction score and is not found in *H. sapiens*. Finally, the MGG suggests the interaction between NUP50-201 and AP2B1. AP2B1 is a subunit of the AP2 complex. AP2 has been predicted to form a complex prior to its nuclear transport [10]. Cytosolic AP2 working with Arkadia regulates endocytosis of EGFR [11], and inhibition of EGFR is a known strategy in lung cancer therapy [12]. However, we were unable to uncover any studies that identify a functional role for AP2 in the nucleus.

##### RRM2-201 contains domain required for membership of Ribonucleotide Reductase

In the MGG model, RRM2-201 is the only RRM2 isoform which contains *ribonuc_red_sm*, the domain required for dimerization and protein interaction with most isoforms of RRM1. Expression of both RRM1 and RRM2 have been implicated in non-small cell lung cancer survival [13] and chemotherapy response [14]. In vivo, the RRM1 and RRM2 form a tetramer structure of two homodimers called the ribonucleotide reductase (RNR). The RNR regulates dNTP levels, which alter DNA replication associated genomic instability [15]. In the survival analysis, a high expression of RRM2-201 was associated with lower survival. Thus, the high expression of RRM2-201 enables RNR membership and subsequent lung cancer susceptibilities. Further, two RRM2B may also be a part of the RNR by replacing the RRM2 homodimer. While the MGG identifies a gained interaction between RRM2-201 and RRM2B, this relationship was only identified using Affinity Capture Mass Spectrometry [16]. There is no evidence to suggest that the interaction exists in vivo or that they may form a dimer inside the same RNR.

##### ASPH-226 is missing active domains

The isoforms of ASPH represent a variety of alternative splicing events and functions. Much of these functions are catalytic, differing based on C-terminal splicing. The ASPH-226 isoform is missing most of the known N-terminal domains and all of the C-terminal domains. Further, ASPH-226 is expressed in each patient, but has a low mean relative expression of 0.023 when compared to other ASPH isoforms. While ASPH expression is associated with lung cancer survival, there is an insufficient amount of annotation of ASPH-226 and its only domain, *Aspartyl beta-hydroxylase N-terminal region*, to identify the mechanism behind its significance. Thus, it is likely that the missing interaction identified in the MGG is only part of the story.
