## Supplementary material for "Uncovering and characterizing splice variants associated with survival in lung cancer patients": Enrichment analysis methods.

### Supplemental material: Methods and results for the enrichment analysis
### Paper: Uncovering and characterizing splice variants associated with survival in lung cancer patients

Sean West[1], Sushil Kumar[2], Surinder K. Batra[2], Hesham Ali[1*], Dario Ghersi[1*]

**1** College of Information and Science Technology, University of Nebraska at Omaha, Omaha, NE, USA **2** Department of Biochemistry and Molecular Biology, University of Nebraska Medical Center, Omaha, NE, USA

*

#### Enrichment of gene lists

We conducted Gene Ontology enrichment on three granularities: (1) the set of survival-significant genes, (2) the genes corresponding to the set of survival-significant splice variants, and (3) the genes corresponding to the splice variants present in all 50 MGGs. Enrichment was conducted using an *in-house* utility called GOUtil. The tool can be found at https://github.com/thecodingdoc/GOUtil and an online version of the enrichment tool is available at funset.uno. This tool uses the hypergeometric test along with FDR $p$-value adjustment. We considered enrichment terms with an FDR $< 0.05$ to be significant. Set subtraction was used to identify enriched terms unique to gene and splice variant granularities.

The union of significant enrichments across the three gene granularities were grouped using spectral clustering, also a part of GOUtil, separately across *biological process*, *cellular component*, and *molecular function*. Representative terms of each cluster were chosen as the medoid terms in each group, as discussed in the documentation for GOUtil. Counts of gene members for each cluster by each granularity were measured.

The three groups were respectively enriched in 77, 159, and 158 GO terms. Unique enrichments were present in each group. We organized enrichment results using spectral clustering. The associated supplemental figure shows the gene counts for each enrichment cluster across each granularity. We note several points of interest in the enrichment results. First, the enrichment of survival-significant genes contained no unique clusters. The gene granularity was not significant to any cellular component terms. Second, the MGGs were enriched in nearly every GO term cluster. Yet, this may be due to the prerequisite of the annotation of MGG genes. In conclusion, the enrichment suggests that the functional information found in gene and splice variant granularities are also found in the MGGs.
