## Supplementary figures and images for "Uncovering and characterizing splice variants associated with survival in lung cancer patients"

### Enrichment term cluster membership across granularities.

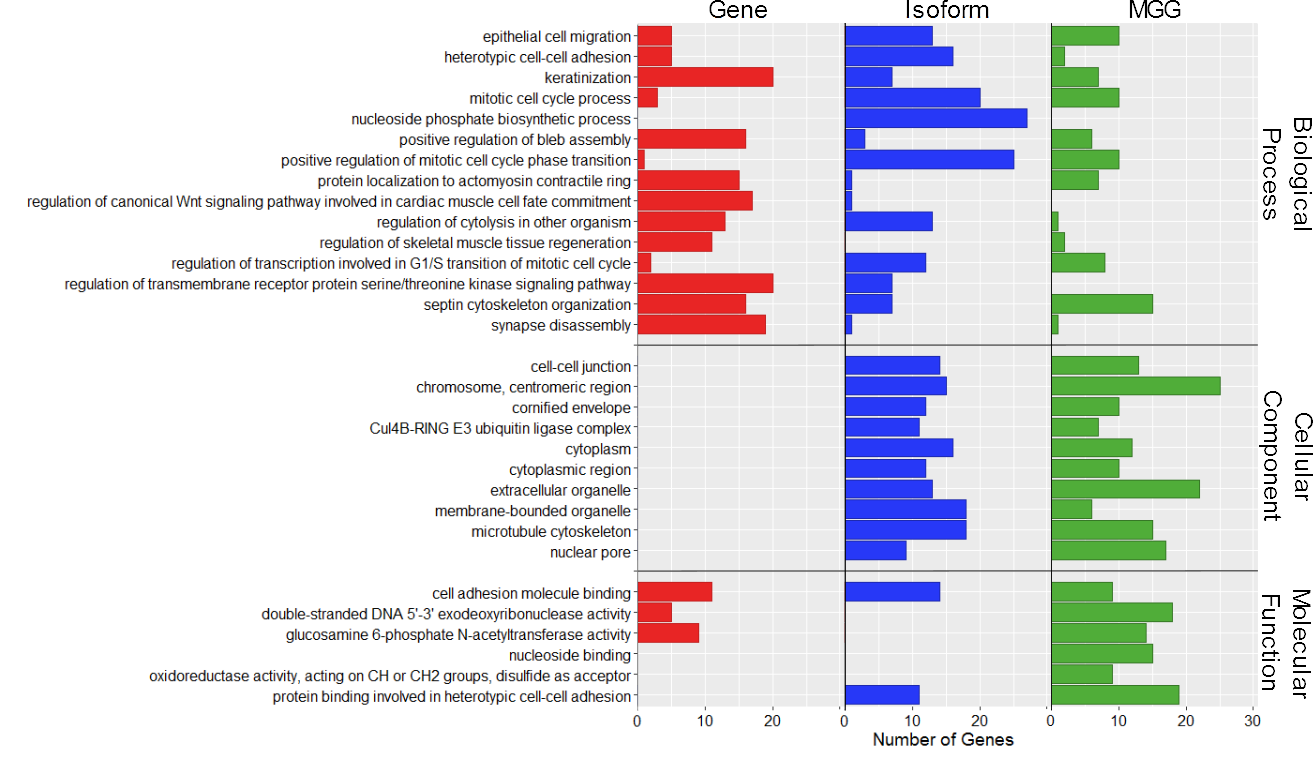
